# Elucidating the role of water in collagen self assembly by isotopically modulating collagen hydration

**DOI:** 10.1101/2023.07.31.551300

**Authors:** Giulia Giubertoni, Liru Feng, Kevin Klein, Guido Giannetti, Yeji Choi, Anouk van der Net, Gerard Castro-Linares, Federico Caporaletti, Dimitra Micha, Johannes Hunger, Antoine Deblais, Daniel Bonn, Andela Šarić, Ioana M. Ilie, Gijsje H. Koenderink, Sander Woutersen

## Abstract

Water is known to play an important role in collagen self assembly, but it is still largely unclear how water-collagen interactions influence the assembly process and determine the fibril network properties. Here, we use the H_2_O/D_2_O isotope effect on the hydrogen-bond strength in water to investigate the role of hydration in collagen self assembly. We dissolve collagen in H_2_O and D_2_O, and compare the growth kinetics and the structure of the collagen assemblies formed in these water isotopomers. Surprisingly, collagen assembly occurs ten times faster in D_2_O than in H_2_O, and collagen in D_2_O self assembles into much thinner fibrils, that form a more inhomogeneous and softer network, with a fourfold reduction in elastic modulus compared to H_2_O. Combining spectroscopic measurements with atomistic simulations, we show that collagen in D_2_O is less hydrated than in H_2_O. This partial dehydration lowers the enthalpic penalty for water removal and reorganization at the collagen-water interface, increasing the self assembly rate and the number of nucleation centers, leading to thinner fibrils and a softer network. Coarse-grained simulations show that the acceleration in the initial nucleation rate can be reproduced by the enhancement of electrostatic interactions, which appear to be crucial in determining the acceleration of the initial nucleation rate. These results show that water acts as a mediator between collagen monomers, by moderating their interactions so as to optimize the assembly process and, thus, the final network properties. We believe that isotopically modulating the hydration of proteins can be a valuable method to investigate the role of water in protein structural dynamics and protein self assembly.

## Introduction

Collagen is the main component of connective tissues such as skin, arteries and bones, imparting to these tissues the mechanical integrity and properties required to ensure their biological functionality(1). The most abundant collagen type in our body is type I. The type I collagen chain contains around 1000 amino acids and is composed of Glycine(Gly)-Xaa-Yaa repeat units, where Xaa-Yaa are often Proline (Pro) and Hydroxy-proline (Hyp). Three polypeptide strands, adopting a left-handed polyproline II-type (PPII) conformation, further associate to form the typical triple helix, tropocollagen(2).Tropocollagens (collagen monomers) self assemble to form intermediate fibrillar structures (microfibrils) that further associate into fibrils with an ordered molecular packing structure that maximizes fibril strength(1). The significance of this packing process is clearly shown in connective-tissue diseases in which it is disturbed due to genetic defects in the collagen type I genes; as a result, the mechanical integrity of diverse tissue types is affected (3).

During the self assembly process, water interacts with and binds to collagen, thereby influencing the mechanical properties of the final network. Water is believed to control collagen properties, for instance, by influencing the structure and stability of the triple helix(4–7) and mediating collagen inter- and intra-molecular interactions(8, 9). In particular, the water layer surrounding the collagen monomers (hydration layer) has been suggested to create a repulsive interaction (“hydration force”) between collagen molecules, arising from the reorganization of the hydration layer that is required for collagen molecules to approach each other closely (9, 10). Research on collagen hydration has focused mostly on the effect of co-solvents, such as ethanol, propanol or glycerol, on the swelling properties of reconstituted fibril films(11, 12). These solvents, however, differ significantly from water, and having very different molecular sizes and dielectric constants than water, they modify not only the collagen hydration but also many other properties. Despite the evident importance of hydration for the properties of collagen fibrils, the mechanism by which water impacts collagen assembly is still largely unclear.

In this work, we study the role of water-collagen interactions on collagen assembly by replacing water with heavy water (D_2_O). The hydrogen bonds between D_2_O molecules are stronger (by ∼10%) than the ones between H_2_O molecules(13, 14). However, contrary to the solvents used previously to investigate collagen hydration, H_2_O and D_2_O have the same electronic structure, and nearly identical molecular size and dielectric constant (78.06 and 78.37, respectively)(15). Hence, changing the isotopic composition of the water can be used to modulate collagen-water interactions, and so study their effect on the assembly process without affecting the electrostatic interactions due to changes in the solvent dielectric constants. A significant effect of D_2_O on protein self assembly has been recently observed for *α*-synuclein (aS) and insulin (INS)(16, 17). In these studies, it was suggested that in D_2_O specific folded structures are stabilized, accelerating (in the case of aS) or slowing down (in the case of INS) the assembly.

Here, we find that the assembly of collagen occurs ten times faster in D_2_O than in H_2_O. This acceleration is somewhat similar to that observed previously for aS(16), but must have a very different origin: collagen has a more stable native ordered structure than aS, and (unlike aS) no drastic refolding of the protein is required for initiating the fibrilization (this refolding being a rate-limiting step for the fibrilization of aS). By combining infrared spectroscopy with atomistic simulations, we find that the faster self assembly observed for collagen in D_2_O is due to the lowering of the energetic penalty of water removal and reorganization at the water-collagen interface, resulting in the enhancement of the initial nucleation rate. Coarse-grained simulations show that the different assembly growth rate and structure in D_2_O can be reproduced by enhancing the electrostatic interactions, which appear to be largely affected by the desolvation energy, and to be central elements into driving the initial nucleation. Our results thus suggest that water guides collagen assembly by slowing down the fibril nucleation by moderating the attractive (mostly electrostatic) interactions between collagen monomers through the creation of a desolvation energy barrier.

## Results

### Network and fibril: kinetics and structure

We first study the influence of the isotopic water composition on the kinetics of collagen self assembly and on the collagen structure at the fibril and network level. We investigate the self assembly kinetics of collagen in H_2_O and D_2_O by using turbidimetry, a standard method (18–21) that relies on the increase in light scattering as the collagen monomers aggregate into fibrils or fibers (inset of Fig.1A). Fig.1A shows the turbidity-time curves measured in heavy water and water at a collagen concentration of 0.1 mg/ml. Both turbidity profiles show the typical sigmoidal growth profile, characterized by a lag phase of near zero turbidity followed by a growth phase with rapidly increasing turbidity. During the lag time (*t*_*lag*_), collagen aggregates grow primarily in length but little in diameter, forming nuclei which have little ability to scatter light. Subsequently, during the growth phase, the collagen monomers anchor onto collagen nuclei, forming fibrils that quickly grow in diameter and molecular weight at a specific growth rate (*k*_*g*_). When the monomers are depleted, the plateau phase is reached (*t*_*plateau*_) as the fibrils attain their mature state(21). The turbidity profiles show that D_2_O samples fibrillate much faster than the H_2_O samples, somewhat similar to *α*-synuclein in D_2_O and H_2_O(16). The *t*_*lag*_ and *t*_*plateau*_ for collagen assembly are ten-fold shorter in H_2_O than in D_2_O, and *k*_*g*_ is one order of magnitude larger in D_2_O with respect to H_2_O. In addition, the final turbidity value, Δ*τ*, is reduced from 0.55 0.07 to 0.39 0.01 in D_2_O, indicating that D_2_O favors the formation of thinner collagen fibrils(22) (this will be further investigated below).

**Fig. 1.**
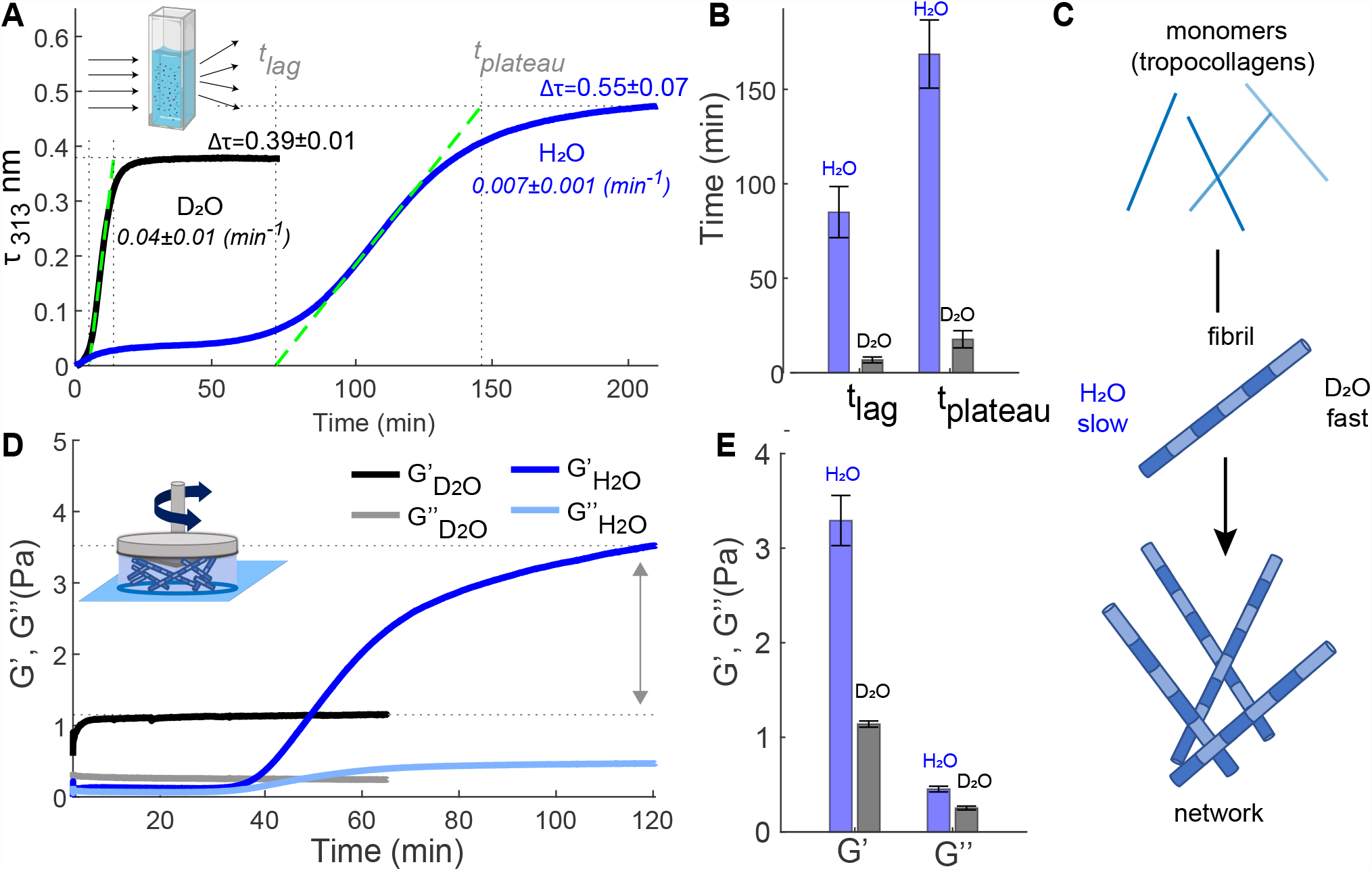
Differences in collagen assembly kinetics and collagen-network elastic properties in H_2_ O and D_2_ O. **A**. Turbidity measurements for water and heavy water solutions containing Type I full-length collagen at a concentration of 0.1 mg*/*ml measured at a temperature of 23°C. Spectra were collected every 15 and 30 seconds for D_2_ O and H_2_ O experiments, respectively. **B**. Lag and plateau time values found for collagen fibrilization in H_2_ O and D_2_ O as described in SI. **C**. Schematic of collagen assembly in water and heavy water. **D**. Rheology measurement for water and heavy water solutions containing collagen at a concentration of 0.5 mg*/*ml. Measurements were conducted at a strain amplitude of 0.8%, an oscillation frequency of 0.5 Hz and temperature of 23°C. **E**. Elastic and viscous moduli after attaining the plateau level.

We then performed rheology measurements to monitor the time-dependence of the mechanical response of the collagen solution during self assembly. The time-dependent elastic and viscous moduli (*G*′ and *G*″, respectively) of a 0.5 mg/ml collagen solution in H_2_O and D_2_O (Fig.1D) show that colla-gen gelates faster in D_2_O as compared to H_2_O, as the elastic modulus reaches its plateau value earlier, consistent with the turbidity measurements. Furthermore, the final elastic modulus in water is ∼400% larger in water than in heavy water, indicating that the network is much softer in D_2_O. Rheological and turbidity experiments in mixed H_2_O:D_2_O (1:1 volume ratio) indicate that D_2_O-induced changes are D_2_O-concentration dependent, with a significant effect already when ∼50% of H_2_O is replaced by D_2_O (see SI, section “HDO Measurements” and Fig. S1-2). Additional frequency-sweep oscillatory rheology measurements reveal that the dynamics of the network relaxation is not influenced by the presence of D_2_O (Fig. S2B).

To investigate the effects of heavy water on the collagen network and fibril, we used confocal microscopy in reflectance mode (CRM) to obtain images of the networks (Fig.2). In water, collagen networks are isotropic and exhibit fan-shaped bundles of fibrils and large pore spaces, similar to the microstructures observed for rat tail Type I collagen in previous studies(22, 23). By contrast, gelation in heavy water does not lead to bundling, and instead a uniform and dense distribution of thin fibers is observed. We then used transmission electron microscopy (TEM) to obtain high-resolution images of the collagen networks (Fig.2B). We find that in H_2_O the network has much larger pore spaces between the collagen fibrils than in D_2_O. Quantitative analysis of the TEM images reveals that the collagen fibrils formed in heavy water are much thinner (Fig.2C): the average fibril thickness is 132±55 nm in H_2_O [in agreement with ref. (22)] and 52±18 nm in D_2_O.

**Fig. 2.**
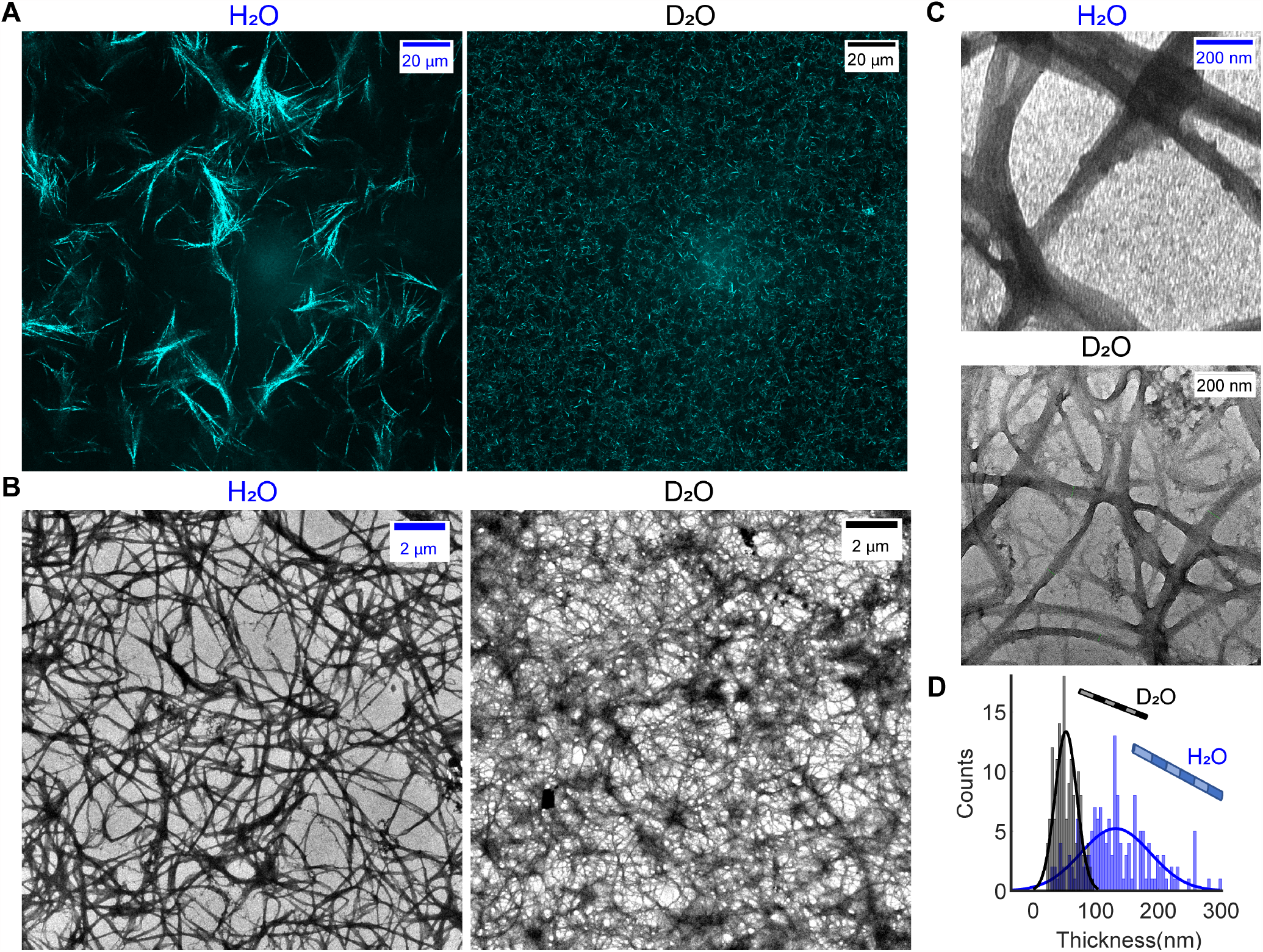
Differences in the collagen fibril and network structures in H_2_ O and D_2_ O. **A**. Representative CRM images of the collagen network formed in water and heavy water at a concentration of 1 mg/ml. **B-C**. Representative TEM images of the collagen network formed in water and heavy water (the images in C are zoomed-in images of B). **D**. Distribution of the fibril thickness in H_2_ O and D_2_ O.

### Water-collagen interactions

To understand the molecular origin of the differences in collagen-assembly kinetics and structure in D_2_O and H_2_O, we compare the structure and hydration of monomeric collagen (its triple-helix structure is shown schematically in Fig.3A) in H_2_O and D_2_O using infrared (IR), two-dimensional IR (2D-IR) and circular dichroism (CD) spectroscopy. IR and 2D-IR spectroscopy probe the local structure and solvation by studying the infrared absorption bands of amide I modes(26–28), whereas CD spectroscopy probes the helicity(29, 30) and the stability(31) of collagen.

**Fig. 3.**
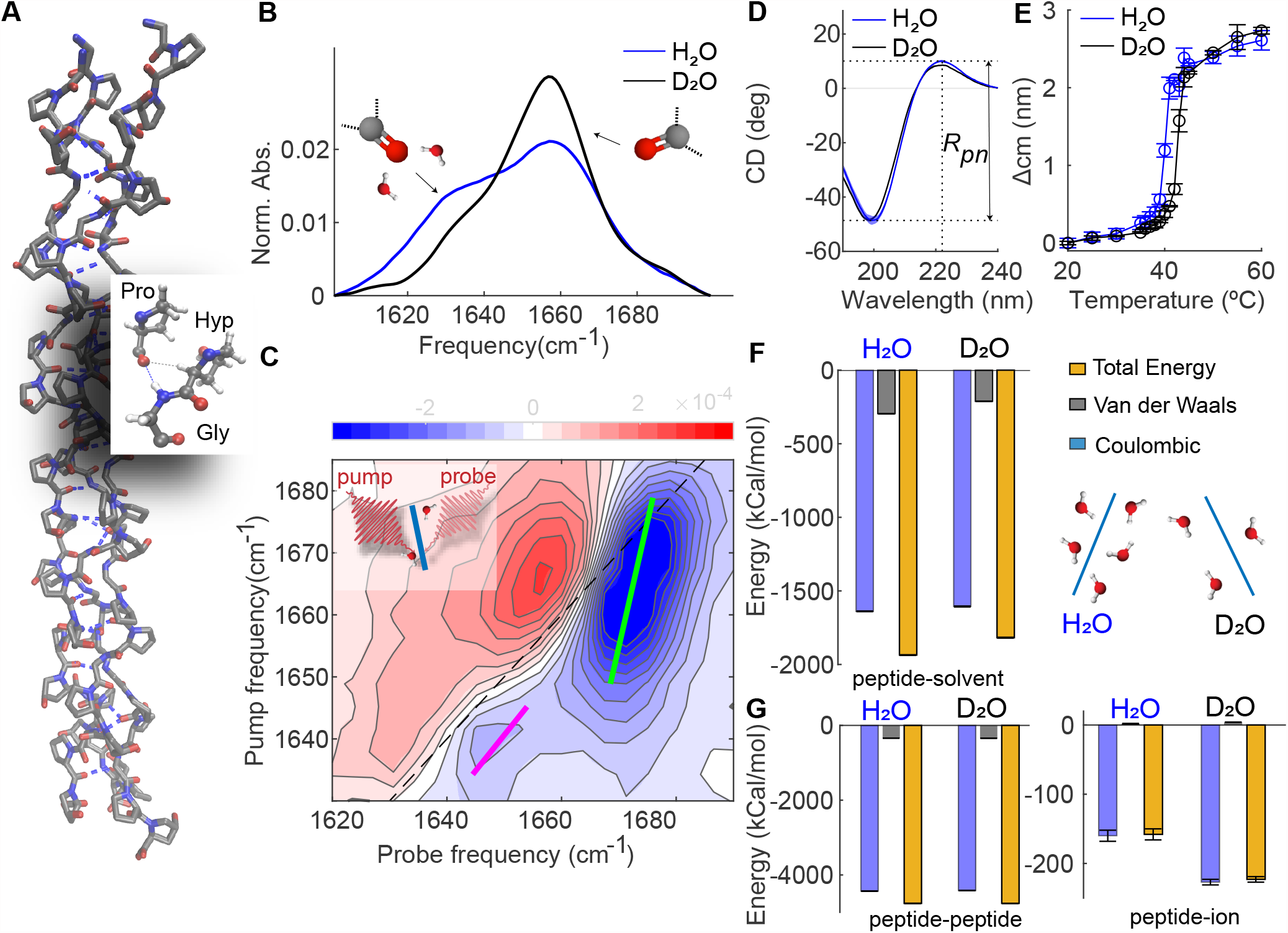
Collagen is less hydrated in D_2_ O than in H_2_ O, but retains the same helicity. **A**. Crystal structure of the (Gly-Pro-Hyp) nonamer (PDB ID: 3B0S(24)). **B**. IR spectra of heavy water and water solutions containing Type I full-length collagen at a concentration of 2 mg/ml and 10 mg/ml, respectively, recorded at 23°C. Full IR spectra are shown in Fig.S3. The IR spectrum in D_2_ O was obtained using FTIR in transmission mode, in H_2_ O it was obtained by using FTIR in reflection mode (ATR-FTIR). In the latter case, because of the shorter optical path length, we use a higher collagen concentration to obtain a sufficient signal-to-noise ratio. The IR spectrum of collagen in water is not concentration dependent (see ref. (25) and Fig.S4). **C**. 2D-IR spectrum of a heavy water solution containing Type I full-length collagen at a concentration of 2 mg*/*ml recorded at a waiting time between pump and probe pulses of 1 ps. The blue contours represent a decrease in absorption (Δ*A <* 0) due to depletion of the *ν* = 0 state, and the red contours an increase in absorption (Δ*A >* 0) due to the induced absorption of the *ν* = 1 → 2 transition. Colored lines in the 2D-IR spectrum represent the calculated central lines (See SI for more details). **D-E**. CD spectra and melting curves extracted from temperature-dependent CD measurements of Type I full-length collagen dissolved in water and heavy water at a concentration of 0.1 mg/ml, respectively (see SI for more details). **F** Interaction energies between the peptide and the solvent (D_2_ O, H_2_ O) molecules and schematic of collagen hydration in D_2_ O and H_2_ O. **G**. Intramolecular energies and energies between the peptide and the ions in D_2_ O and H_2_ O.

In Fig.3B, we report the normalized IR spectra of the triple-helical collagen monomer dissolved in D_2_O and H_2_O recorded at a temperature of 23 °C. We observe two main bands at 1635 cm^−1^ and 1660 cm^−1^, in agreement with literature(25, 32–34) with the 1660 cm^-1^ band more intense in heavy water with respect to water. The low frequency band was previously assigned to the vibrations of carbonyl that are accessible and well-exposed to water(34); to verify this assignment we use 2D-IR spectroscopy, a technique that can provide direct information on the collagen hydration(27). In pump-probe 2D-IR spectroscopy, we use a tunable narrowband pump pulse to excite molecular vibrations at a specific frequency *ν*_pump_, and measure the pump-induced change in absorption Δ*A* at all frequencies using a broad-band probing pulse. Each vibrational mode of a molecule gives rise to a +*/* − doublet on the diagonal(27). In the 2D-IR spectrum reported in Fig.3C, we observe two pairs of diagonal peaks at pump frequencies of 1635 cm^-1^ and 1660 cm^-1^. The lineshapes of the two diagonal peaks differ significantly, with the 1660 cm^-1^ diagonal peak being more tilted with respect to the diagonal. The dependence of the 2D-IR response on the pump frequency is a measure of the inhomogeneous broadening of the IR band(27), which is due to a distribution of transition frequencies caused by solvent-protein interactions. The degree of inhomogeneity can be characterized by calculating the inverse value of the slope of the 2D-IR bleaches (central line slope or CLS)(35); and we find that the CLS values for the 1635 and for the 1660 cm^-1^ peaks are 0.7±0.15 and 0.35±0.11, respectively (values and errors represent the mean and standard errors obtained over 3 different measurements). The higher CLS value for the band at low frequency indicates a larger inhomogeneity, and thus a broader frequency distribution, than for the peak at 1660 cm^-1^. The broader frequency distribution is due to interactions between functional groups and solvent molecules, indicating that the amide groups absorbing at 1635 cm^-1^ experience better solvation than the ones absorbing at 1660 cm^-1^. We then fit the IR spectra in D_2_O and H_2_O (Fig.3B) by using Gaussian-shaped peaks (see SI and Fig.S4C-D for more details). We find that the area of the peak of the well solvent-exposed carbonyl decreases by ∼30% in intensity when collagen is dissolved in D_2_O as compared to H_2_O. This spectral difference [also observed in ref.(25)] was found to be independent of collagen concentration and amide H/D exchange (Fig. S4D-F). Furthermore, an increase in the ratio between less- and well-solvated carbonyl bands is observed in collagen fibril solutions when the fibrillation takes place in D_2_O (Fig S4A); but also in collagen dissolved in H_2_O (with the temperature set to 4°C to prevent fibrillation)(33).

We investigated whether the reduced hydration in D_2_O influences the helicity of the collagen triple helix using CD spectroscopy. Fig.3D shows the CD spectra of collagen dissolved in water and heavy water at a concentration of 0.1 mg/ml. Both CD spectra have a minimum at 198 nm and a maximum at 220 nm, the typical spectral signatures of the collagen triple-helix.(29) To check whether the reduced solvation affects the collagen helicity, we calculate the ratio between the intensities of the maximum and the absolute of the minimum values, *R*_*pn*_ (an experimental criterion for triple-helicity)(30). We find identical ratios (∼0.19) in H_2_O and D_2_O, indicating a similar helicity of collagen. In addition, we extracted the melting temperatures of the collagen triple helix from the temperature dependence of the CD spectra (see Fig.3E and SI for more details), resulting in 40±1°C and 43±1°C in H_2_O and D_2_O, respectively. This result indicates that the collagen monomer has a less stable structure in H_2_O than in D_2_O, in agreement with previous studies on collagen-based peptides(31).

To validate the experimentally observed reduction in water-collagen interactions in D_2_O compared to H_2_O, we performed molecular dynamics simulations of the (Gly-Pro-Hyp) nonamer triple helix starting from its crystal structure (PDB ID: 3B0S(24), Fig.3A). Two sets of simulations were carried out in H_2_O and D_2_O at 300 K. In each case, five independent copies were run for a cumulated sampling time of 10 *µ*s. The triple helices are structurally stable over the course of the simulations with an average root-mean-square deviation from the crystal structure of 2.4 ±0.002 Å. The energetic analysis reveals that the total interaction energies between the nonamer and the solvent are less favorable in D_2_O than in H_2_O (−1818 ±3 kcal/mol in D_2_O and − 1936±2 kcal/mol in H_2_O, Fig.3F). The reduction of water-protein interaction is also reflected in the smaller number of solvent-collagen hydrogen bonds in deuterated water (131±0.4) with respect in water (136± 0.4), which confirms the experimentally observed reduced hydration in D_2_O, and complements previous results on other biomolecules(36, 37). The total number of the total intramolecular energies(Fig.3G) and intramolecular hydrogen bonds are essentially identical, while the interaction with the ions is more favorable in D_2_O (− 223±4 kcal/mol) than in H_2_O (− 158 ± 8 kcal/mol). The latter is probably an effect of the reduced hydration in D_2_O as compared to H_2_O. Thus, the molecular dynamics simulations show a reduction in water-collagen interactions in D_2_O, leading to a less solvent-exposed and more stable protein structure, as we observed in IR and 2D-IR measurements, without significantly altering the collagen helicity, as we observed in the CD measurements.

### Collagen-collagen interactions

How can partial dehydration of collagen in D_2_O modify the collagen-collagen interaction in such a way as to cause the observed changes in assembly kinetics and structure? To address this question, we perform coarse-grained molecular dynamics simulations using collagen-mimetic molecules. The assembly of collagen is known to be driven and regulated by an interplay between hydrophobic and electrostatic interactions(11, 38–45), and coarse-grained simulations have proven successful in revealing how this interplay controls the self assembly(46). To see which of these forces is most strongly influenced by the reduced hydration in D_2_O, we systematically modify them in the simulations and see if we can reproduce the experimentally observed changes in collagen assembly rate and fibril structure. In the coarse-grained MD simulations, collagen molecules are described as elastic rods that carry a pattern of charges (Fig.4A). The rods can interact with each other via screened electrostatic interactions as well as via generic, hydrophobic attractions and have previously been shown to form clusters and collagen-like fibrils (46). Electrostatic interactions are modeled using a Debye-Hückel potential, while a Lennard-Jones potential is used for hydrophobic interactions. To study how the reduced hydration in D_2_O can cause the observed changes in assembly rate and fibril structure, we vary the strength of electrostatic and hydrophobic interactions, and average the obtained results over ten independent simulation runs for each set of parameters. Snapshots of these simulations are shown in Fig.4B. The results (Fig.4C-D) show that increasing the hydrophobic interaction strength decreases the assembly rate and increases fibril diameter, the opposite of the experimentally observed trend. However, upon increasing the electrostatic interaction strength, the assembly rate is increased and fibril diameter is decreased, exactly as is observed experimentally in D_2_O (Figs. 1 and 2). These results indicate that the experimentally observed acceleration of assembly as well as the thinner fibrils in D_2_O compared to H_2_O can be effectively reproduced by enhancing electrostatic interactions, rather than by enhancing hydrophobic interactions.

**Fig. 4.**
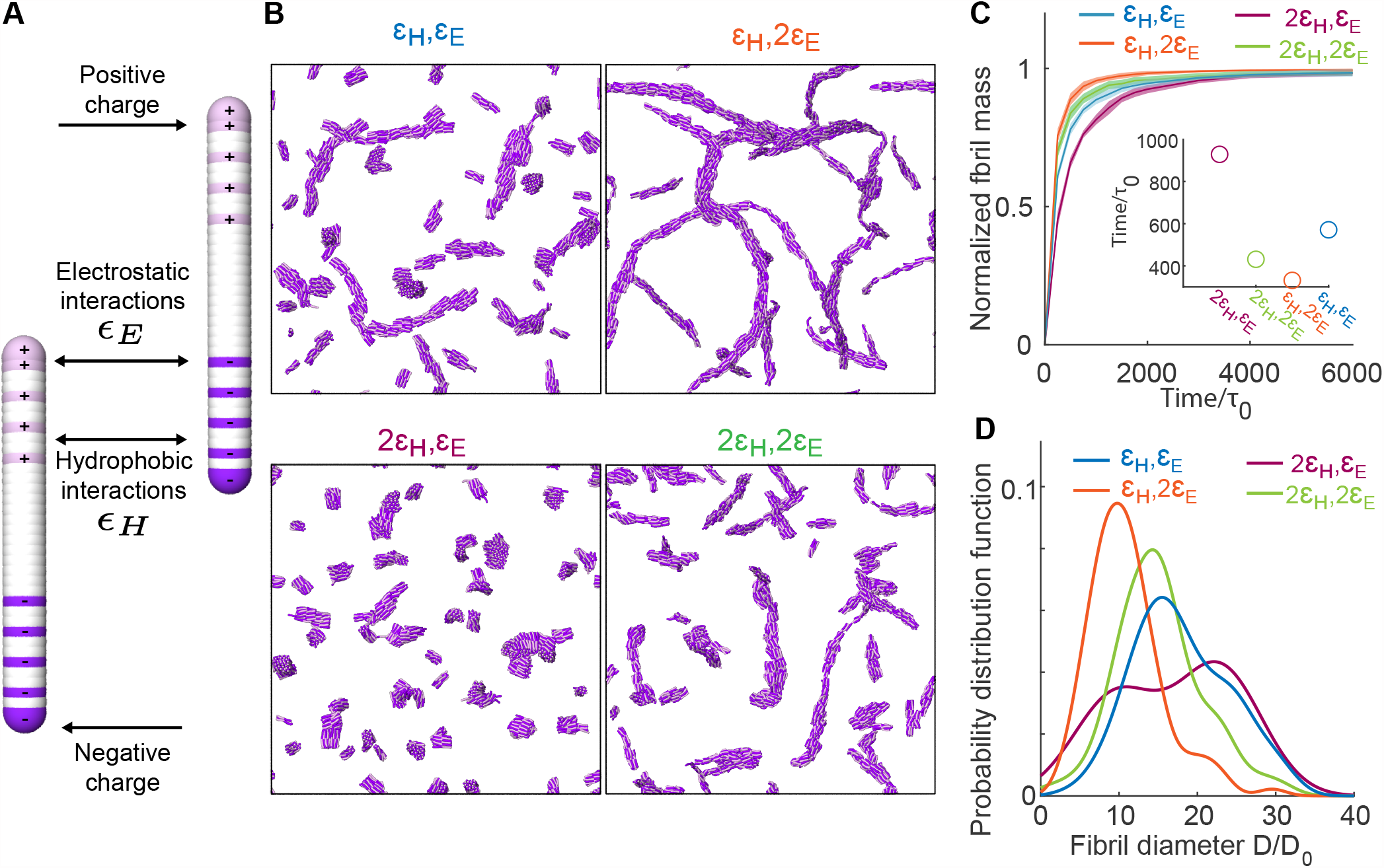
Coarse-grained simulations can qualitatively reproduce the differences in collagen fibril and network structure in H_2_ O and D_2_ O. **A**. The collagen-mimetic molecules are simulated as elastic rods made of overlapping beads. The beads carry charges as indicated (positive are pink, negative are purple, and white is neutral). On top of these electrostatic interactions, all the beads between different molecules interact via generic hydrophobic interactions. **B**. Simulation snapshots of the equilibrated system for different combinations of electrostatic and hydrophobic interactions (ϵ _*H*_ = 0.05*kT*, ϵ_*E*_ = 5*kT*). **C**. Normalized fibril mass as a function of simulation time. The inset shows the time at which the assembled mass reaches 80% of the total monomer mass, where *τ*_0_ is the MD unit of time. **D**. Probability distribution function of the fibril diameter *D* normalized by the smallest measured fibril diameter *D*_0_.

## Discussion

Our results show that changing the solvent from H_2_O to D_2_O causes a ten-fold acceleration of the collagen assembly (Fig. 1), and a dramatic structural change and softening of the final fibril network (Fig. 2). These differences become with increasing D_2_O concentration, with a significant effect on the kinetics and mechanical properties already at a 1:1 ratio of H_2_O:D_2_O (Fig. S1-2). In the nucleation-growth mechanism of collagen self assembly, the fibril diameter is determined mainly during the nucleation step(47). Our coarse-grained simulations suggest that in D_2_O more nucleation centers form due to the enhancement of electrostatic interactions between collagen monomers (Fig. 4). These nucleation centers compete with each other for the remaining collagen monomers, so that the increased attractive interaction in D_2_O results in the thinner fibrils that are observed in the TEM images and the coarse-grained simulations. This is consistent with the thinner collagen fibrils formed at higher temperature, which also accelerates fibrilization(23). By combining (2D-)IR with CD and MD simulations we find that water-collagen interactions are reduced in D_2_O, leading to a more stable and less water-bound structure, without altering the collagen helicity. These findings are consistent with previous studies which have shown that D_2_O is a poorer protein solvent than H_2_O, and that D_2_O favors a less water-exposed but more stable protein structure(31, 48–57), affecting the assembly properties (16, 17, 58).

How can a reduction in collagen hydration affect the assembly process and the fibril structure so dramatically? For proteins to interact, the water molecules, which are tightly bound to the hydrophilic groups on the protein surface, have to be removed and reorganized, leading to a large energetic penalty (desolvation energy) for the protein assembly. Recently, it has been suggested that this desolvation energy plays a crucial role in the assembly of intrinsically-disordered proteins (IDP)(59): although the assembly into fibrils is thermodynamically favoured by the entropic gain in solvent release, the fibril nucleation is limited by the large desolvation energy. The initial self assembly rate can thus be increased in less hydrating conditions, resulting in a faster assembly. Our results indicate that the reduced hydration in D_2_O affects the assembly process (and final fibril and network properties) in a similar manner, by lowering the desolvation energy barrier, which limits the initial nucleation. The coarse-grained simulations show that the acceleration in the initial nucleation rate can be reproduced by enhancement of the electrostatic interactions, suggesting that these interactions (rather than hydrophobic ones)are more sensitive to changes in the desolvation energy, and play a more important role in driving the initial nucleation. Although we believe that further experiments should be carried out to fully understand this last aspect, the role of hydrating water molecules in modulating electrostatic interactions has already reported for other proteins(59, 60), and a similar enhancement of the attractive interaction between collagen monomers as a result of a lower desolvation energy in D_2_O seems plausible.

In earlier studies, it was already suggested that a short-range repulsive “hydration force” might be crucial for the structure and properties of collagen fibrils, and that the penalty associated to restructuring the tightly bound water molecules might prevent collagen molecules from coming too close to each other (9, 10, 12). However, so far this potential role of desolvation energy in collagen assembly was only indicated indirectly in experiments and simulations (11, 40, 41). The unique possibility of isotopically modulating the hydration while keeping the other solvent properties the same makes it possible to directly demonstrate the crucial role of the desolvation in collagen assembly. Our results indicate that water controls the mechanics of collagen networks by moderating attractive interactions between collagen monomers that guide the self assembly. In this way, water drives the formation of few initial nuclei rather than many competing ones, ensuring the cooperative nature of collagen self assembly. In the future, it would be interesting to determine whether specific regions in the collagen sequence, and if so which ones, are most important in establishing this mediating role of water. Furthermore, by exploiting the different hydration in D_2_O and H_2_O, we intend to probe this mediating role for collagen interactions with other tissue components, such as minerals in bones(61) and hyaluronic acid in cartilage(62).

Our findings provide new insights into how hydration modulates collagen properties to finely tune the mechanics of living tissues(63), and suggest new avenues toward the design of artificial collagen-based materials. Controllable and tunable macroscopic properties might be achieved by subtle changes in the solvent isotopic composition instead of alterating the chemical structure of a biomaterial’s building blocks. Finally, altered water-collagen interactions are believed to play a role in several age-related diseases(6, 64–66), and to partially contribute to tissue dysfunction in these disorders. It is well-known that genetic defects in the collagen type I genes COL1A1 and COL1A2 can cause osteogenesis imperfecta, Caffey disease and Ehlers-Danlos syndrome with a distinct bone or skin pathology, but our limited knowledge of the collagen folding hierarchy and its tissue-specific interfering factors set us still far from comprehending the mechanisms leading to such hyperostosis or fragility of bones, skin or blood vessels(67–69). The results presented here show that collagen hydration modulates the assembly rate and diameter of fibrils, which are also impacted in these diseases(70–72). It is therefore not unlikely that modified hydration may exacerbate the molecular defects of collagen type I (i.e. excessive posttranslational modification, misfolding) in determining the phenotypic outcome. We hope that further studies will give insight in the way that water distribution influences collagen quality, and how this might potentially be used for therapeutic purposes(73).

## Conclusion

We have shown that changing the solvent from H_2_O to D_2_O induces a tenfold acceleration of collagen assembly, and leads to thinner fibrils and a much softer collagen network, with significant effects already observable when 50% of H_2_O is replaced by D_2_O. By combining spectroscopy with molecular dynamics simulations, we have found that collagen in D_2_O is less hydrated than in H_2_O, and that it adopts a less water-exposed and more stable structure without altering its helicity. Our results indicate that the kinetic and structural changes originate from a lower energetic penalty for water removal and water reorganization at the collagen surface in D_2_O, and coarse-grained simulations suggest that this desolvation energy influences mostly electrostatic interactions, which appear to be crucial in determining the nucleation rate. Our results directly demonstrate the role of hydration in collagen self assembly: the water layer surrounding the collagen acts as a mediator, moderating collagen-collagen interactions in order to slow down the assembly so as to optimize the final network properties.

## Material and Methods

### Sample preparation

Lyophilized collagen containing telopeptides (Type I collagen from rat tail tendon, Roche cat. no. 11179179001) was purchased from Sigma Aldrich. Deuterated materials used are: D_2_O (deuterium oxide; Sigma-Aldrich, no. 151882-25G); Acetic acid-d_4_ (Sigma-Aldrich, no. 233315-5G); NaOD (sodium deuteroxide; Sigma-Aldrich, no. 372072-10G). Collagen was dissolved in water, heavy water or 1:1 water:heavy water solutions containing 0.1 or 0.2% (v/v) of acetic acid (Emsure, no. 1000632500). The collagen was dissolved to obtain a final concentration of 0.2 mg/ml (for turbidity/CD measurements), 1 mg/ml (for rheology measurements) and 2 mg/ml or 10 mg/ml (for (2D-)IR measurements in heavy water and water, respectively), and stored at 4°C for at least 4 days before usage ensuring collagen is fully dissolved. All collagen samples were prepared on ice to prevent early self assembly and the self assembly was initiated by neutralizing acidic collagen solutions. First, weighing collagen in an Eppendorf tube and subsequently adding an equal volume of customized buffer solution to obtained a final pH of 7.3-7.5 and ionic strength I = 0.17 M. The customized buffer solution is made of Milli-Q water, 10× PBS (made with phosphate buffered saline tablet; VWR, no. E404-100TABS) solution, and 0.1 M NaOH (made with 1M sodium hydroxide solution; Honeywell Fluka, no. 319511-500ML). The customized buffer solution contained a fraction of 20% of PBS, whereas the Milli-Q:NaOH is 60%:20% or 40%:40%, depending on the initial acetic acid concentration. The pH of all the collagen solutions were measured by using a pH-meter (Thermo Scientific, Orion 2 Star) that was calibrated for measuring the pH in H_2_O solutions instead of D_2_O solutions. The measured pH* of a D_2_O solution was transformed to the pH value by using the following equation: *pH* = (*pH*\* + 0.4) 0.929 (74).

### Infrared Spectroscopy

For IR-measurements of heavy water solutions, sample containing collagen at a concentration of 2 mg/ml was placed in a circular sample cell composed by two CaF_2_ windows separated by a 100 *µ*m spacer. Measurements were done in transmission mode using a Bruker Vertex 70. Per measurement, 32 scans were made, with a spectral resolution of 2 cm^-1^. The temperature was kept at 23°C using a temperature controller (Julabo, TopTech F32-ME). The frequency range was from 7000 cm^-1^ to 400 cm^-1^. For IR-measurements of water solutions, sample containing collagen at a concentration of 10 mg/ml (or 5 mg/ml) was measured in reflection mode using a PerkinElmer Frontier FT-IR spectrometer fitted with a Pike GladiATR module equipped with a diamond ATR-crystal (ø = 3 mm). Spectra were averaged over 20 scans. Temperature was maintained at room temperature by using a built-in heating/cooling plate. The spectrum of the solvent was subtracted to obtain the individual spectrum of the collagen.

### Two-dimensional infrared spectroscopy

A detailed description of the setup used to measure the 2DIR spectra can be found in ref. 75. Briefly, pulses of wavelength 800 nm and with a 40 fs duration are generated by using a Ti:sapphire oscillator, and further amplified by using a Ti:sapphire regenerative amplifier to obtain 800 nm pulses at 1 kHz repetition rate. These pulses are converted in an optical parametric amplifier to obtain mid-IR pulses (∼ 20 *µ*J, ∼ 6100 nm) that has a spectral full width at half max (FWHM) of 150 cm^−1^. The beam is split into a probe and reference beam (each 5%), and a pump beam (90%) that is aligned through a Fabry-Pérot interferometer. The pump and probe beams are overlapped in the sample in an ∼250-*µ*m focus. The transmitted spectra of the probe (*T*) and reference (*T*_0_) beams with pump on and off are recorded after dispersion by an Oriel MS260i spectrograph (Newport, Irvine, CA) onto a 2×32-pixel mercury cadmium telluride (MCT) array. The probe spectrum is normalized to the reference spectrum to compensate for pulse-to-pulse energy fluctuations. The 2DIR signal is obtained by subtracting the probe absorptions in the presence and absence of the pump pulse.

### Circular dichroism

CD spectra were recorded with a JASCO CD spectrometer (Model: J-1500-150) in the far-UV at wavelengths, *λ*, ranging from 180 nm to 260 nm to obtain information on the secondary structure of the proteins. Data were recorded with a data pitch of 0.2 nm, a scan speed of 20 nm/min, a digital integration time of 0.5 s, and an optical path length of 1 mm. Spectra were smoothened using the Savitzky-Golay filter built-in in the spectrometer software. Temperature-dependent measurements were performed at temperatures ranging from 20 °C to 60 °C at increments of 5 °C with an equilibration time of 4 min. At 35-45 °C smaller increments of 1 °C were used with 8 min equilibration time. From each experiment the spectrum of the buffer was subtracted and the results of the three experiments were averaged for the final analysis.

### Turbidity

The kinetics study of collagen self assembly was performed on a UV-Vis spectrophotometer (Agilent Technologies, Cary 8453). Neutralized cold collagen solutions were pipetted into plastic cuvettes (Brand, UV-Cuvette micro, no. 759220), which were quickly sealed with a cover to avoid evaporation and H/D isotopic exchange and subsequently placed in the water-jacked cuvette holder. Measurements were performed at 23 °C. Spectra were recorded every 15 s and 60 s after neutralization for heavy water and water samples, respectively. The spectrum of the respective solvent was used as a background. As collagen self assembly proceeded, the absorbance at a wavelength of 313 nm (*A*_313_) were recorded as a function of time. Increase of *A*_313_ over time during collagen self assembly represents an increase in scattering. The absorbance readings were converted into turbidity values (*τ*) by using the relation: *τ* = *A*_313_ *ln*10.

### Rheology

Rheology study of collagen was performed with a stress-controlled rheometer (Anton Paar, Physica MCR 302), equipped with a cone-plate geometry (50 mm diameter, 1° cone angle, 100 *µ*m gap). The bottom plate temperature was controlled using a Peltier element. Neutralized cold collagen solutions at a concentration of 1.25 mg/ml (experiments shown in SI) or 0.5 mg/ml (experiments shown in the main text) were pipetted onto the plate, and the cone was immediately lowered to the measuring position. We used a thin layer of low viscosity mineral oil (Sigma-Aldrich, no. 330760-1L) around the sample to prevent solvent evaporation and H/D isotopic exchange. Within ∼2 min the oscillatory rheology measurement was started.

### CRM

To prepare collagen samples for CRM measurements, we used the protocol described in ref.(62) Briefly, neutralized cold collagen solution was pipetted into the customized sample holder, composed of two coverslips and the adhesive silicone isolator (Thermo Fisher Scientific, Press-to-Seal silicone isolator) in between. The coverslips were cleaned beforehand with isopropanol and Milli-Q water and dried by nitrogen flushing. The sample holder was then immediately placed into a petri dish and sealed by parafilm to prevent solvent evaporation and H/D isotopic exchange. Collagen at a concentration of 1 mg/ ml was left to polymerize at 23°C. Both water and heavy water samples were measured after at least 150 min at 23°C from neutralization to attain full network formation. The equilibrium collagen network images were taken by an inverted confocal laser scanning microscope (Leica Stellaris 8 platform) equipped with a 63x, NA = 1.30 glycerol-immersion objective (Leica), a (supercontinuum) white light laser with laser line 488 nm for illumination and the reflected light was dectected with silicon multi-pixel photon counter (Leica, Power HyD-S) detector. Glycerol (Leica, ISO 836) was used for objective immersion.

### TEM

To prepare collagen samples for TEM measurements, we used the protocol described in ref.(76). Briefly, after neutralization, fibril assembly was initiated by placing the samples in a closed container (comprised of the cap of a closed Eppendorf tube placed upside down) for at least 150 minutes. The collagen fibrils were transferred to glow-discharged electron microscopy grid by peeling off the collagen gel drop surface with the grid (purchased from QuantiFoil, C support Cu400), which was left on the collagen surface between 1 to 12 hours. The sample was then washed 1 or times by placing a drop of milliQ water and blotting the drop without completely drying the grid. Finally, the sample was stained by adding a drop of 2% uranyl acetate and blotting it to dryness. TEM images were analyzed by using ImageJ, which is an image analysis and open source software(77). After scale calibration, thickness of the fibrils was calculated by taking the width of different fibrils in at least 4 different images of 4 different samples for a total of 150 measured thickness points. Width measurements were taken from the non-smoothed image by manually drawing a line perpendicular to the long axis of the bundle or the filament between the edges of the fibril. The edges were determined as the location where the darkened region produced by the defocus halo starts.

### Molecular dynamics simulations

The crystal structure of collagen-mimetic peptides composed of showing nine Gly-Pro-Hyp repeats (PDB ID: 3B0S (24)) was used as starting conformation. Two sets of simulations of the collagen triple helix {Gly-Pro-Hyp}_9_ were carried out in water and heavy water at 300 K, cumulating 10 *µ*s (5×2*µ*s runs). All simulations were run using the GROMACS 2020.4 software package (78, 79), the CHARMM36m forcefield (80) and explicit solvent molecules, *i*.*e*. TIP3P for water and modified TIP3P-HW for heavy water (81).

Each collagen triple helix was solvated in a cubic box (12 nm per edge), with TIP3P (82) or TIP3P-HW (81) water molecules, to which 140 mM NaCl were added to mimic experimental conditions. The N- and C-termini were uncapped. Periodic boundary conditions were applied and the time step was fixed to 2 fs. Following the steepest descent minimization, the system were first equilibrated under constant pressure for 5 ns, with position restraints applied on the heavy atoms of the protein, followed by 5 ns NPT equilibration in absence of restraints. The temperature and the pressure were mantained constant at 300 K and 1 atm, respectively by using the modified Berendsen thermostat (0.1 ps coupling) (83) and Berendsen barostat (2 ps coupling) (84). The production simulations were performed in the NVT ensemble in absence of restraints. The short range interactions were cut-off beyond distances of 1.2 nm, and the potential smoothly decays to zero using the Verlet cut-off scheme. The Particle Mesh Ewald (PME) technique (85) was employed (cubic interpolation order, real space cut-off of 1.2 nm and grid spacing of 0.16 nm) to compute the long range electrostatic interactions.

### Coarse grained molecular simulations

Our model is based on the “D-mimetic” molecule, which is a synthetic collagen-mimetic molecule, that has been shown to self-assemble into collagen-like fibrils (46, 86). Since this D-mimetic protein consists of 36 amino acids only, our molecule consists of 36 beads that are arranged into a linear chain. With *σ* being the MD unit of length, each of these beads measures *r* = 1.12*σ* in diameter and is in contact with its direct neighbors via a harmonic bond *E* = *κ*_bond_(*r*−*r*_0_)^2^, where *κ*_bond_ = 500*kT /σ*^2^ is the bond strength and *r*_0_ = 0.255*σ* is the equilibrium distance. This results in a molecule length of *l* = 10*σ* and consequently, *σ* = 1 nm, because the D-mimetic peptide has a length of 10 nm. We use an angular potential *E* = *κ*_angle_(*θ* −*θ*_0_)^2^ that acts between three neighboring beads to define the rigidity of our molecule, where *κ*_angle_ = 50*kT* controls the molecular rigidity and *θ*_0_ = *π* is the equilibrium angle. Additionally, all beads carry a unit charge with respect to the charge distribution of the D-mimetic molecule, as shown in Fig. 4A in the main text. All the beads on different molecules are able to interact with each other via a generic, hydrophobic potential described by a cut-and-shifted Lennard-Jones potential 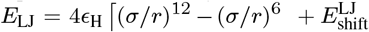, if two interacting beads are at a distance *r < r*_*c*_ = 2*σ*, and is 0 otherwise, *ϵ*_H_ is the strength of nonspecific or hydrophobic interactions, which is one of our control parameters. Furthermore, two charged beads *i, j* are able to interact with each other via a cut-and-shifted screened electrostatic potential (DLVO) 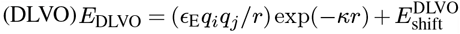 a distance *r < r*_*c*_ = 2*σ*, and 0 otherwise, *κ* = 1*σ* is the screening length and its length of 1 nm corresponds to the Debye screening length at physiological conditions. *ϵ*_E_ defines the effective strength of the electrostatic interactions and is the second control parameter we will explore, while *q*_*i*_ represents the sign of the charge of bead *i* (*q*_*i*_ = 1). Since neighboring beads in a molecule have overlapping volume and distances between charges in the same molecule can be small, we exclude interactions of beads in the same molecule for 1-2, 1-3, 1-4, and 1-5 neighbors. The simulations are initialized by randomizing the positions and orientations of *N* = 2500 molecules in a cubic box of length *L* = 171*σ*, resulting in a molecule number concentration of *c*_mol_ = 0.0005*σ*^−3^. We integrate the system in the NVE ensemble (V, E being the volume of the box and the total energy of the system, respectively) with a Langevin thermostat to simulate Brownian motion of the molecules, with the LAMMPS MD package (87). Our integration timestep is 0.001*τ*_0_, where *τ*_0_ denotes the MD unit of time, and the damping coefficient was chosen to be 1*τ*_0_.

## ACKNOWLEDGEMENTS

We thank Dr. Steven Roeters (Aarhus University), Dr. Federica Burla, Prof. Dr. Nico Sommerdijk (Radboud Medical Center, Nijmegen, The Netherlands) and Prof.Dr. Mischa Bonn (Institute for Polymer Research, Mainz, Germany) for the useful discussion. We thank Dr. Wim Roeterdink and Michiel Hilberts for the technical support. GK acknowledges financial support by the “BaSyC - Building a Synthetic Cell” Gravitation grant (024.003.019) of The Netherlands Ministry of Education, Culture and Science (OCW) and The Netherlands Organization for Scientific Research and from NWO grant OCENW.GROOT.2019.022. This publication is part of the project (with Project Number VI.Veni.212.240) of the research programme NWO Talent Programme Veni 2021, which is financed by the Dutch Research Council (NWO). We acknowledge support from the Sectorplan Bèta & Techniek of the Dutch Government.

