## Supplementary for "Elucidating the role of water in collagen self assembly by isotopically modulating collagen hydration"

### Supplementary materials for: Elucidating the role of water in collagen self assembly by isotopically modulating collagen hydration

**Giulia Giubertoni[1,✉], Liru Feng[1], Kevin Klein[4,8], Guido Giannetti[1], Yeji Choi[5], Anouk van der Net[3], Gerard Castro-Linares[3], Federico Caporaletti[1,2], Dimitra Micha[7], Johannes Hunger[5], Antoine Deblais[2], Daniel Bonn[2], Andela Šarić[4], Ioana M. Ilie[1,6], Gijsje H. Koenderink[3], and Sander Woutersen[1]**

[1]Van 't Hoff Institute for Molecular Sciences, University of Amsterdam, Amsterdam, The Netherlands
[2]Van der Waals-Zeeman Institute, Institute of Physics, University of Amsterdam, Amsterdam, The Netherlands
[3]Department of Bionanoscience, Kavli Institute of Nanoscience Delft, Delft University of Technology, Delft, The Netherlands
[4]Institute of Science and Technology Austria, Klosterneuburg, Austria
[5]Max Planck Institute for Polymer Research, Mainz, Germany
[6]Amsterdam Center for Multiscale Modeling (ACMM), University of Amsterdam, the Netherlands
[7]Amsterdam University Medical Centers (UMC), Vrije Universiteit Amsterdam, the Netherlands
[8]UCL, London, United Kingdom

DRAFT

**A. Turbidity curve analysis.** To identify the rate of increase ($k$) of the turbidity curves, we need to find the time range where the growth is linear. To do so, we used the first-derivative curves, based on a similar approach used in ref.(1). We selected the time range between the time value where the first derivative is 75% of the maximum derivative, and then we fitted a line using least squares fitting. The slope of the fitted line is $k$. The $t_{lag}$ is calculated by taking the time at which the tangent to the turbidity curve at the maximum slope is 0, whereas $t_{plateau}$ is calculated by taking the time at which the tangent is equal to the maximum turbidity value.

**B. HDO measurements.** We studied collagen assembly and properties in HDO solutions, containing a $H_2O$: $D_2O$ at a volume ratio of 1:1. In Fig.1, we reported the turbidity measurements performed at 1 mg/ml in $H_2O$and HDO. For experiments in HDO, we used the stock solution of collagen dissolved in water containing 0.2 wt % of acetic acid (same stock solution of the $H_2O$ experiments); and we then neutralized it by using a customized $D_2O$-buffer to obtain a final collagen concentration of 1 mg/ ml with a 1:1 volume ratio of $H_2O$:$D_2O$. The reported continuous curves and shades represented the mean and standard deviation over three different measurements. We noticed that assembly in HDO happens faster than in $H_2O$. To check whether there is a dependence on the solvent used for the stock solutions, we repeated the same measurements by using the stock solution of collagen dissolved in heavy water containing 0.2 wt % of acetic acid, and neutralizing it with customized $H_2O$-buffer to obtain again a final collagen concentration of 1 mg/ ml with a 1:1 volume ratio of $H_2O$:$D_2O$ ( dashed grey line in Fig.1). We again observed a faster assembly than in $H_2O$, indicating an independence on the solvent used for the original stock solution. In Fig.2A, we reported the rheological measurements performed at 1.25 mg/ml of collagen in $H_2O$, HDO and $D_2O$. In this case for the HDO measurements, we used a stock solution of collagen dissolved in HDO containg 0.2 wt % of acetic acid. Similar to the turbidity measurements, we observed a faster fibrillization in HDO; further, the elastic modulus was lower in HDO compared to $H_2O$, indicating a significant effect on the collagen gel properties already when 50% of water was replaced with heavy water. Additionally, we performed frequency-sweep (10-0.1 Hz) oscillatory rheology measurements of equilibrium collagen network with a concentration of 1.25 mg/mL of collagen concentration at a strain of 0.5% (Fig.2B). The presence of $D_2O$ doesn't influence the dynamics of stress relaxation in the collagen network With a power-law with an exponent of $\sim$0.1 for all collagen networks, G' is weakly dependent on the frequency, $\omega$. This is typical behavior of biopolymer networks that are transiently cross-linked by non-covalent interactions.(2, 3)

**C. Fit analysis for IR spectroscopy.** The linear absorption spectra (Fig. 3B of the main text) were fitted using 3 Gaussian-shaped peaks absorbing at 1635, 1660, and 1680 $cm^{-1}$. The fits were performed by leaving the width and center frequencies of the peaks as free parameters. As shown in Fig. 4B-C, the fits well reproduce the linear infrared data.

**D. Central line slope (CLS) for 2D-IR spectroscopy.** To calculate the CLS, we first fit a series of cuts through the 2D spectrum that are parallel to the probe frequency axis by using two Lorentzian-shaped peaks that describe the bleach (blue color) and the excited state absorption (red color). We then fit a line through the center positions of the bleach obtained from the fit of the cuts. The line slope is then the CLS value.

**E. Melting temperature calculation.** The CD spectra for solutions of collagen at ambient temperatures exhibited a marked negative peak at $\sim$198 nm and an adjacent positive peak at $\sim$220 nm (Fig.S5). With increasing temperature both collagen in $H_2O$ and $D_2O$denature, as suggested by the decrease of the amplitude of the negative 200 nm peak and the change of the sign of the 220 nm spectral feature (Fig.S5A-B). These transitions were observed in a very narrow temperature range of around 39-40°C for solutions of collagen in $H_2O$ and around 41-43°C for solutions of collagen in $D_2O$(Fig.S5A-B). These different ranges indicated a higher melting temperature of collagen in $D_2O$, as compared to $H_2O$. To quantify solvent isotope effects on the melting behavior of collagen, we calculated the center of mass wavelength of the negative 220 nm peak:

$$CM = \frac{\sum_{\lambda=180\ nm}^{210\ nm} \theta_{CD} \bullet \lambda}{\sum_{\lambda=180\ nm}^{210\ nm} \theta_{CD}} \tag{1}$$

The center of mass wavelength, CM, was obtained as the mean value of the wavelength($\lambda$), weighted by the absolute value of the ellipticity ($\theta_{CD}$) at 180-210 nm. The determined values for the three individual scans for collagen in $H_2O$ and $D_2O$ are displayed in Fig.S5C, where we observed that the transition temperature of collagen in $D_2O$ shifts to higher temperatures, as compared to collagen in $H_2O$. To quantify the transition temperature, we averaged the data of the 3 repeat measurements per condition in Figure 2 and fitted a sigmoidal function together with a linear variation of the CM as a function of temperature $T$ to the data:

$$CM\,(T) = a_1 + \frac{a_2 - a_1}{1 + 10^{(a_3 - T)}a_4} + a_5 T \tag{2}$$

where $a_j \;\; (j = 1 \cdots 5)$ are fit parameters. As displayed in Fig.S5C, Eq (2) describes the data very well, and the obtained parameters are listed in table 1. From these fits we found the melting temperature ($a_3$) to increase from 40.2 °C in $H_2O$ to 42.8 °C in $D_2O$.

DRAFT

**F. Analysis of coarse-grained simulations.** In order to quantitatively analyze our simulations, we recorded the positions of all molecules every 10000 timesteps for a total of $6 \cdot 10^6$ timesteps, resulting in a total simulation time of $6000\tau_0$ and 600 simulation snapshots (frames). To measure the assembly rate, in each frame we calculated the mass of all assembled clusters normalized by the total mass, where a cluster was defined as an aggregation of $n \geq 10$ monomers. To calculate the fibril diameter probability distribution function, we identified fibrils in the last frame of a simulation as clusters that contained at least $n \geq 10$ monomers and were at least $l \geq 2l_{\mathrm{monomer}}$ long. The diameter $D$ of such a fibril was calculated as the average diameter along its longitudinal axis normalized by the smallest measured diameter $D_0$ among all identified fibrils. To obtain averages and standard deviations for the assembled mass and to increase the number of fibrils available for the diameter distribution, we ran 10 independent simulations for each set of parameters $(\epsilon_{\mathrm{H}}, \epsilon_{\mathrm{E}})$.


DRAFT

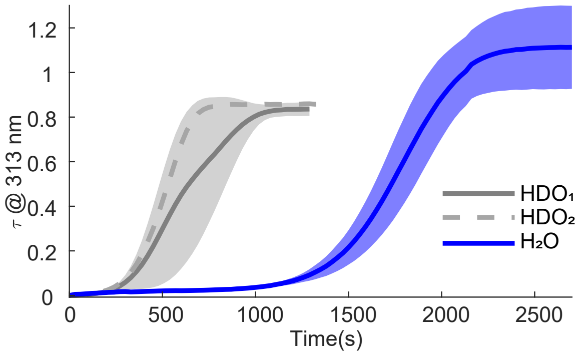


**Fig. 1.** Turbidity curves for collagen fibril formation in HDO and $H_2O$ at concentration of 1 mg/ml at 23°C. Stocks solutions of collagen dissolved in water or heavy water (0.2 wt % of acetic acid, pH=3.3=3.4) were neutralized using heavy water buffer or water buffer to obtain a 1:1 ratio of $H_2O$( which we refer to as $HDO_2$) and $D_2O$ ( which we refer to as $HDO_1$). Shaded areas represent the standard deviation obtained over 4 and 3 different measurements for HDO and $H_2O$,respectively.

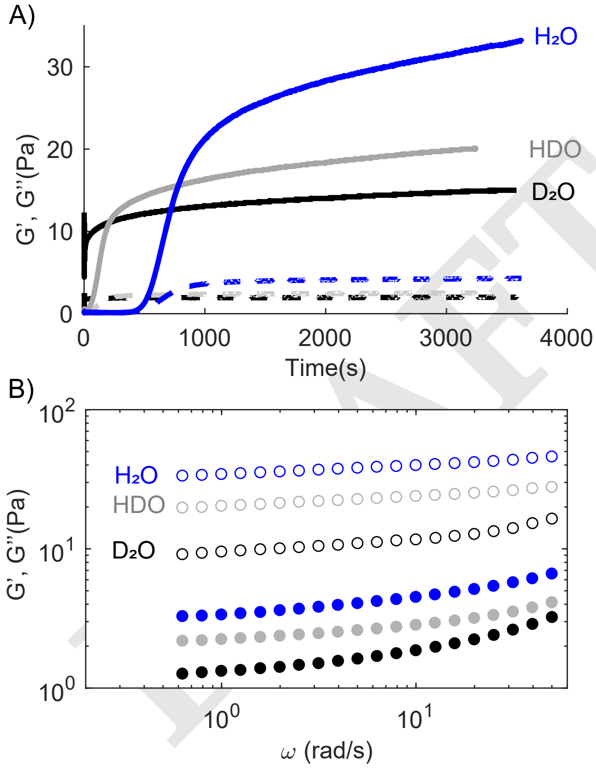


**Fig. 2.** A) Representative evolution of elastic shear modulus G' (continuous lines) and viscous shear modulus G" (dashed lines) during collagen self-assembly in $D_2O$, HDO and $H_2O$. Measurements were conducted with a collagen concentration of 1.25 mg/mL at a strain amplitude of 0.5%, a frequency oscillation of 0.5 Hz and temperature of 23°C. B) Frequency sweep of fully formed collagen network in different solvents with a collagen concentration of 1.25 mg/mL at a strain amplitude of 0.5% and temperature of 23°C. (G' = opened symbols,G"= çlosed symbols ).

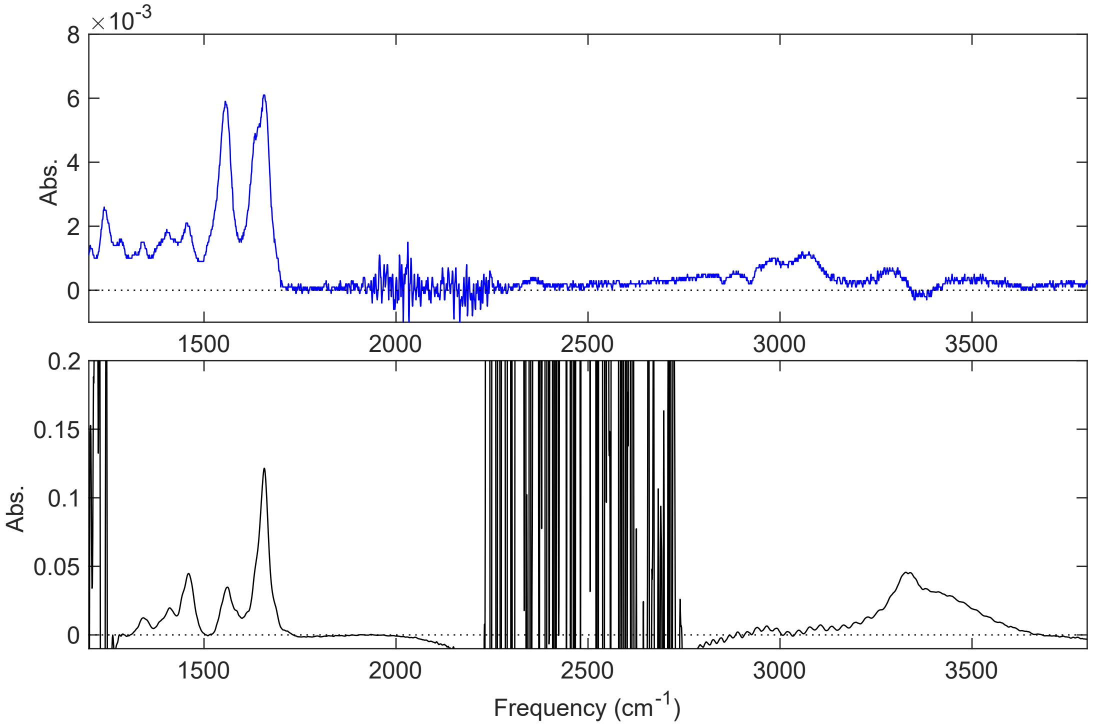


**Fig. 3.** IR spectra of collagen monomers dissolved in water and heavy water ( 0.2 wt% acetic acid, pH=3.2-3.4) recorded between 1200 and 3600 $cm^{-1}$ (top and bottom respectively). Amide II vibration (O=C-N-H) absorbs at around 1550 $cm^{-1}$, whereas Amide II' (O=C-N-D) absorbs at 1490 $cm^{-1}$.

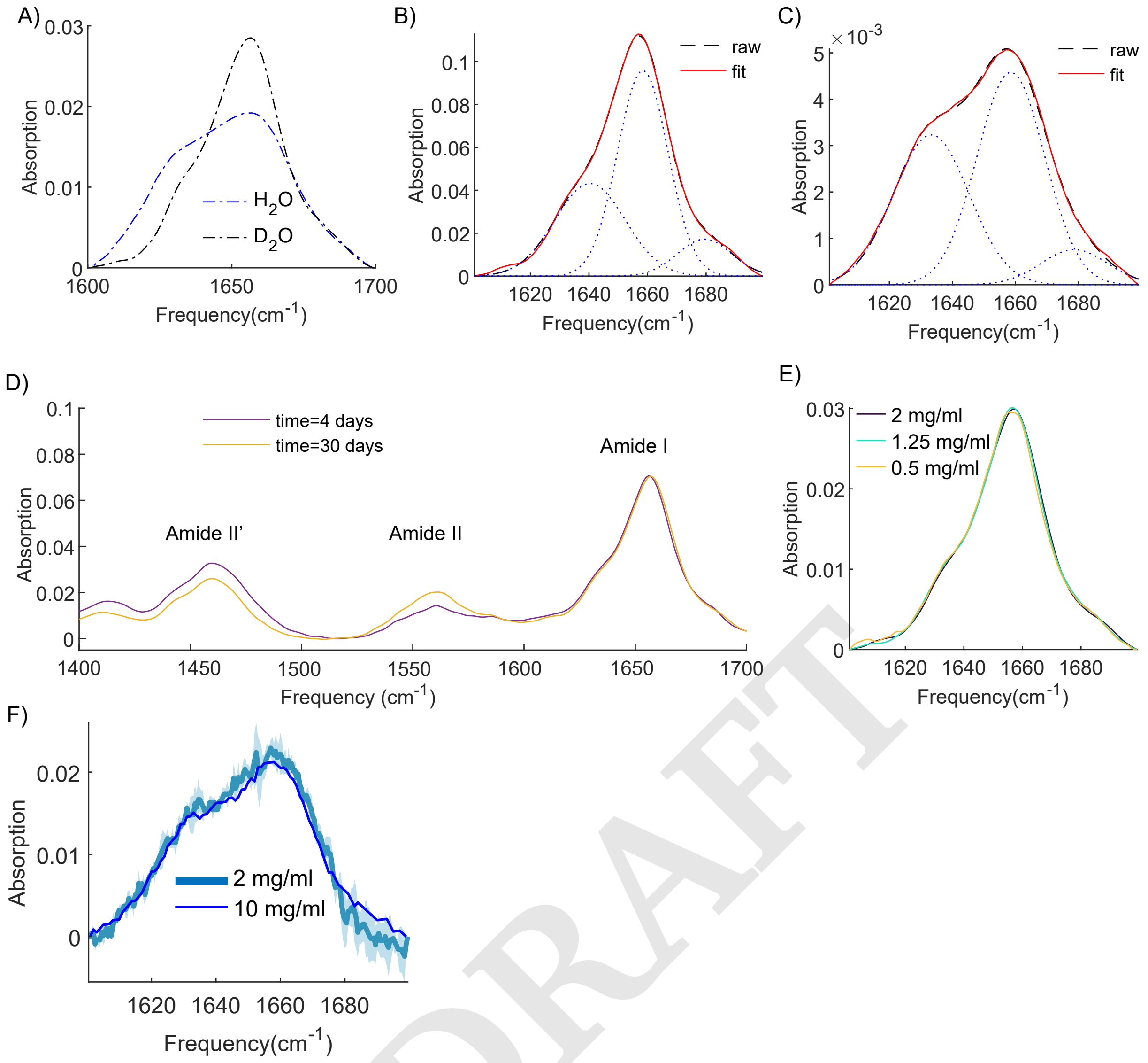


**Fig. 4.** A. IR spectra of collagen fibrils dissolved in water and heavy water(pH=7.4) at a concentration of 5 mg/ml and 1.25 mg/ml, respectively. B.-C. Fit of the IR spectra of collagen monomers dissolved in water and heavy water ( 0.2 wt% acetic acid, pH=3.2-3.4) at a concentration of 10 mg/ml and 2 mg/ml, respectively. Blue dotted lines represent the fitted Gaussian-shaped peaks, and the red line the fit to the raw data (black line). D. IR spectra of collagen at different level of N-H/N-D exchange, which slowly increase over storage time. Upon N-H/N-D exchange, the amide II frequency shifts from 1550 $cm^{-1}$ to 1490 $cm^{-1}$, whereas the amide I frequencies red-shift of 1-2 $cm^{-1}$. We observe that the ratio between the peak intensities of the two amide I bands is constant, although the amide II decreases in intensity because of the N-H to N-D exchange. By using collagen samples using the same stock solution after different storage time, we found no evidence of N-H/N-D effect on the macro- and microscopic changes observed when dissolving collagen in heavy water. E-F. IR spectra of collagen monomers dissolved in heavy water and water ( 0.2 wt% acetic acid, pH=3.2-3.4) at different concentration. The light blue line and shade represent the mean and the standard deviation obtained from 3 different measurements. All measurements reported here were done at 23 °C.

DRAFT

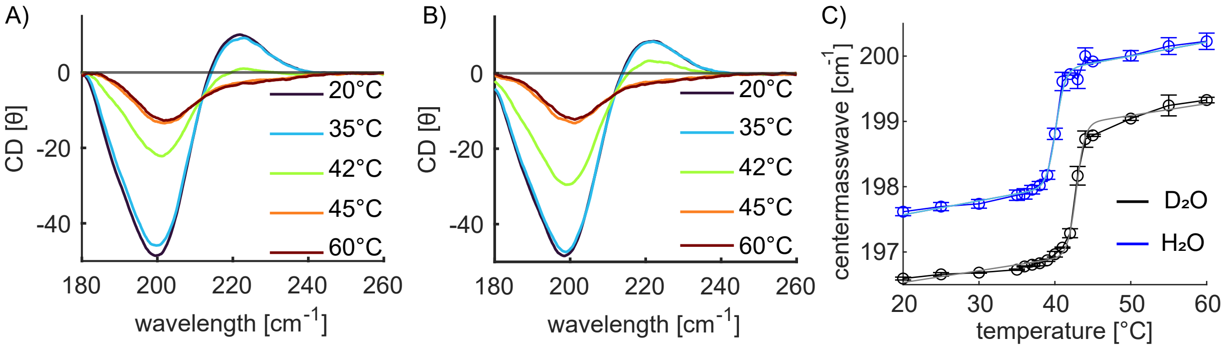


**Fig. 5.** A.-B. CD spectra of solutions of collagen monomers dissolved in water (A) and heavy water (B) ( 0.2 wt% acetic acid, pH=3.2-3.4) at a concentration of 0.1 mg/ml at different temperatures. C. Averaged center of mass wavelength as a function of temperature. Symbols show experimental data and error bars show the standard deviation within three experiments. Solid shaded lines show fits of Eq.(2) to the data, while solid opaque lines are guides-to-eye.
